# Contractile ring mechanosensation and its anillin-dependent tuning during early embryogenesis

**DOI:** 10.1101/2022.09.09.506560

**Authors:** Christina Rou Hsu, Gaganpreet Sangha, Wayne Fan, Joey Zheng, Kenji Sugioka

**Affiliations:** Life Sciences Institute, The University of British Columbia, 2350 Health Sciences Mall, Vancouver, BC V6T1Z3, Canada; Department of Zoology, The University of British Columbia, 2350 Health Sciences Mall, Vancouver, BC V6T1Z3, Canada

**Author notes:** **Corresponding author** Kenji Sugioka, Life Sciences Institute, Department of Zoology, The University of British Columbia, 2350 Health Sciences Mall, Vancouver, BC V6T1Z3, Canada. These authors are equally contributed (Christina Rou Hsu, Gaganpreet Sangha).

## Abstract

The contractile ring plays crucial roles in animal morphogenesis. Previous studies have explored how tissue mechanics controls the contractile ring; however, the mechanisms by which the ring senses tissue mechanics remain largely unknown. Here, we demonstrate the mechanism of contractile ring mechanosensation and its tuning during asymmetric ring closure of *Caenorhabditis elegans* embryos. High-resolution imaging of cortical flow revealed that local suppression of the ring-directed cortical flow is associated with a delay in furrowing. This suppression of cortical flow results from cortical compression. We found that the artificial inhibition of ring-directed cortical flow was sufficient to induce asymmetric ring closure in symmetrically dividing cells. Moreover, genetic analysis suggests that the positive feedback loop among ring-directed cortical flow, myosin enrichment, and ring constriction constitutes the anillin-dependent, mechanosensitive engine driving asymmetric ring closure. Our results suggest that the balance between RhoA-dependent and cortical flow-dependent myosin enrichment fine-tunes the ring’s mechanosensitivity in tissues.

## Introduction

The cytokinetic contractile ring plays crucial roles in morphogenesis during asymmetric cell division and epithelial morphogenesis. Previous studies have shown that intracellular and extracellular mechanics can influence contractile ring positioning, but how the contractile ring senses these mechanical cues remains largely unknown ^1–3^. One important model system to study contractile ring mechanosensation is the asymmetric ring closure found in animal zygotes and epithelial tissues, termed unilateral or asymmetric cytokinesis ^4, 5^. Unilateral cytokinesis is ubiquitously observed in both invertebrates and vertebrates, including humans ^6–10^, and its dysregulation has been associated with reduced cytokinesis resilience, disrupted epithelial integrity, and defective lumen morphogenesis^6, 11, 12^.

Contractile ring assembly is regulated by the activation of small GTPase RhoA. The mitotic spindle plays a central role in specifying the site of RhoA activation through the centralspindlin complex ^13–16^. The activated RhoA then induces Rho-associated kinase (ROCK)-dependent myosin activation, as well as formin-dependent actin assembly, to initiate contractile ring formation ^17–19^. Furthermore, the centrosome and astral microtubules at the polar region suppress ring assembly ^20–27^. Pioneering studies have shown that asymmetric spindle positioning is sufficient to induce asymmetric ring closure in a few examples such as ctenophore zygotes and normally symmetrically-dividing sea urchin embryos, ^28, 29^ and that the spindle-cortex proximity asymmetrically activates RhoA ^30^.

Conversely, unilateral cytokinesis in various vertebrate and invertebrate epithelial tissues and in the *C. elegans* zygote is spindle positioning-independent. In both systems, the mere spindle-cortex proximity does not dictate the leading edge of the ring^6, 31–34^. In *Drosophila* embryonic and follicular epithelia, adherens junctions regulate asymmetric ring closure ^2, 34^. In this system, passive mechanical resistance imposed by adherens junctions anchors the contractile ring at the apical cell cortex. In contrast, unilateral cytokinesis in the *C. elegans* zygote is regulated by contractile ring components anillin and septin^6^. In this system, contractile ring components enrich at the leading edge, forming a structurally asymmetric ring. However, the precise mechanisms of asymmetric ring closure remain unclear in both systems. In *C. elegans* zygotes, knockdowns of anillin and septin disrupt the asymmetric ring closure ^6^; however, the *ani-1(RNAi)* phenotype is rescued in a mutant background of a worm-specific RhoA activator, NOP-1 ^35^. Furthermore, anillin and septin are dispensable for asymmetric ring closure at least in *Drosophila* embryonic and adult dorsal thorax epithelial cells, leaving the role of anillin cryptic ^2, 36^.

Recent studies show that cortical flow, a concerted flow of cell cortex materials at the cell surface, facilitates contractile ring assembly through cortical compression^37–39^. Therefore, we hypothesized that the contractile ring utilizes this mechanism to sense tissue mechanics. To elucidate the mechanism of contractile ring mechanosensation, we performed high-resolution 4D imaging. By developing analysis pipelines measuring various parameters such as ring closure, ring eccentricity, cortical flow, and myosin enrichment rate from the same datasets, we gained an integrative view of mechanochemical contractile ring regulation during asymmetric ring closure. Quantitative analysis of these parameters suggests that the furrowing delay at the lagging cell cortex is induced by the mechanical retardation of cortical flow towards the ring. By conducting an *in vitro* cell-bead adhesion assay, we artificially reduced the ring-directed cortical flow, thereby confirmed the role of ring-directed cortical flow in asymmetric contractile ring closure. Finally, through a genetic analysis, we identified the molecular pathway that tunes contractile ring mechanosensitivity.

## Results

### High-resolution 4D imaging and quantitative analysis of contractile ring dynamics

To better understand the mechanism of asymmetric ring closure, we performed high-resolution 4D imaging of GFP-fused non-muscle myosin II (NMY-2) in *C. elegans* zygotes (Figures 1A and S1A). We applied a new imaging method combining the use of spacer beads and a refractive index-matching sample medium to avoid the previously reported compression-dependent cell rotation ^40, 41^ and to improve image resolution ^42^, respectively (see Methods for detail). These improvements allowed us to capture uncompressed embryonic volumes (30 µm thickness) at 5.6-s intervals with a 1 µm step size, without cell rotation (Figure S1B and Movie S1).

**Figure 1.**
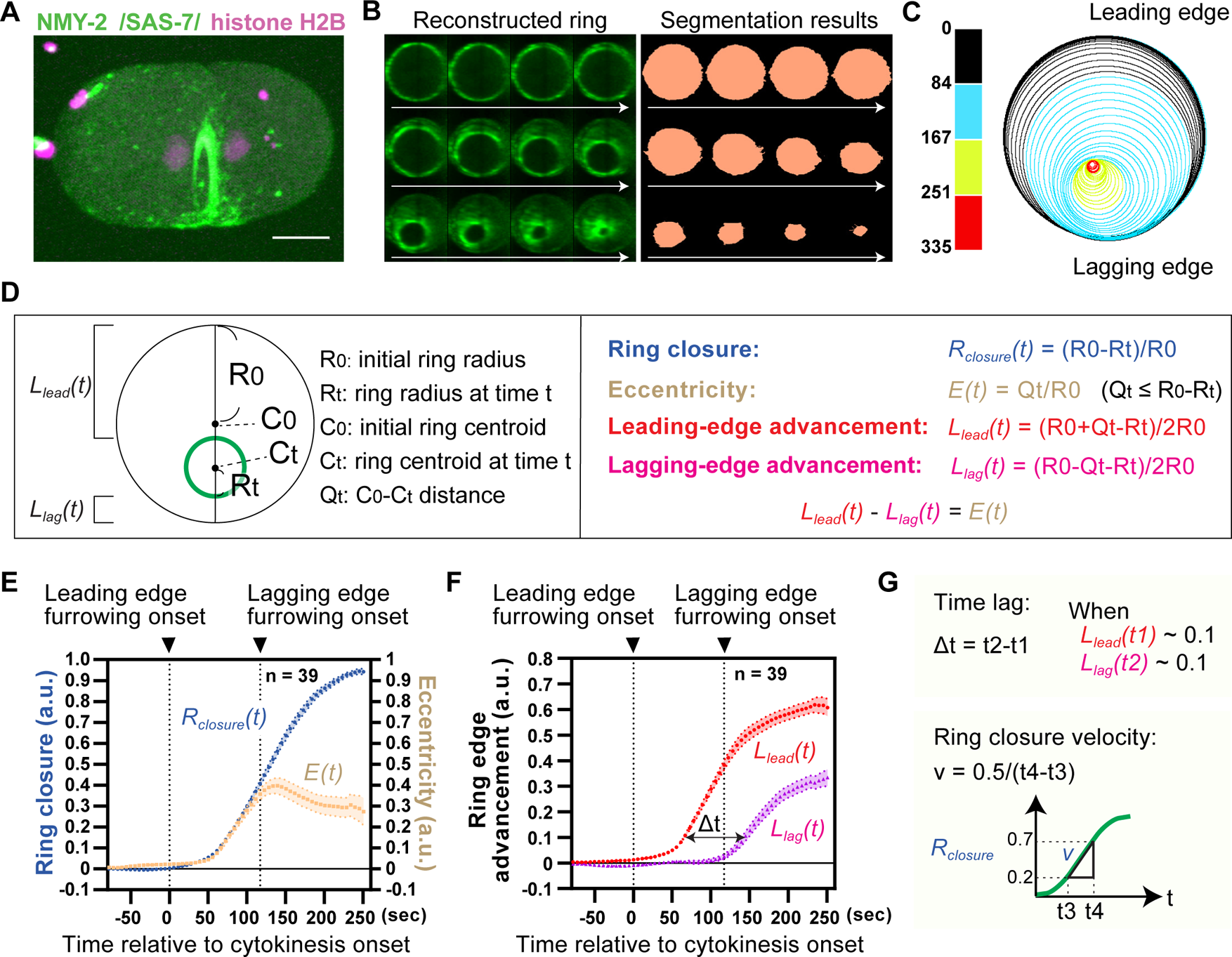
Quantitative analysis of asymmetric contractile ring closure in the *C. elegans* zygote. (**A**) Asymmetric contractile ring closure in *C. elegans* zygote. NMY-2::GFP (green, non-muscle myosin II), GFP::SAS-7 (green, centriole) and histone H2B::mCherry (magenta, chromosome) are shown. In this and subsequent figures, anterior and posterior are to the left and right, respectively. (**B**) Computationally reconstructed ring and segmentation results. The top left image shows cytokinesis onset, and the second and third rows highlight the advancement of the leading and lagging ring edges, respectively. See Fig S1 for method and full frame images. (**C**) Trajectory of ring constriction. Outlines of the contractile ring during cytokinesis are plotted based on the segmentation results. The color coding indicates time relative to cytokinesis onset. (**D**) Geometric analysis of the contractile ring closure. Eccentricity and ring edge advancement are the degree of ring off-centering and the distance between the ring edge and initial ring perimeter, respectively. (**E**) Mean ring closure and eccentricity. (**F**) Mean ring edge advancement. Lagging furrowing onset was defined as the timing when the L_lag_(t) exceeded 0.02 (2% relative to initial ring size). (**G**) Definitions of time lag and ring closure velocity (See text). Times are relative to cytokinesis onset. Error bands indicate 95% confidence intervals. Scale bars, 10 µm.

We developed image analysis pipelines to quantify the dynamics of contractile ring closure (Figures 1B and S1A). First, we performed image segmentation to computationally detect the contractile ring (Figures 1B and S1C). Using the segmentation data, we measured the ring radius and ring center coordinates at every time point. From these measurements, we calculated the ring closure indices *R_closure_(t)* (with 1.0 representing complete closure), ring eccentricities *E(t)* (indicating off-center tendency), and the degrees of leading-edge and lagging-edge advancement towards the cell center *L_lead_(t)*, *L_lag_(t)* (Figure 1D). Due to the natural variations in cytokinesis timing, we aligned the time series data relative to the time around 10% contractile ring closure (Figure S1D). We also defined cytokinesis onset based on the extrapolation of an initial linear part of the ring closure curve to 0 (Figure S1D). Markedly, the newly developed imaging and quantification methods allowed us to detect differences in sub-percentage ring closure indices. The data obtained show that the contractile ring initially undergoes nearly the upper limit of eccentric closure (Figure 1E; Movie S1), and unilateral furrow ingression continues until the start of lagging edge furrowing (Figure 1E–F). These data indicate that the primary factor contributing to ring off-centering is the time interval between the onsets of furrowing at the leading edge and the lagging edge (Figure 1F; 0-117 sec).

### Ring closure velocity and a delay in lagging edge furrowing determine the ring eccentricity

We analyzed the relationship among peak eccentricity, time lag between leading and lagging edge advancement, and ring closure velocity. The time lag, Δt, was determined by the delay between the leading and lagging edges reaching 10% advancement relative to the initial ring diameter (Figure 1F–G). Additionally, ring closure velocity, *v*, was calculated based on the slope of the R_clousre_ curve between 20% and 70% closure points (Figure 1G). We found that ring closure velocity and peak eccentricity have no clear correlation (Pearson’s r = 0.352, *p* = 0.028, n = 39; Figure 2A). On the contrary, the time lag is weakly correlated with the peak eccentricity (Pearson’s r = 0.54, *p* = 0.0003, n = 39; Figure 2A). Since a slowly closing ring might have a longer time lag than a faster one, we calculated the normalized time lag, Δt_n_, by multiplying the ring closure velocity by the time lag (Figure 2B). We found that the normalized time lag is strongly correlated with peak eccentricity (Pearson’s r = 0.93, *p* < 0.0001, n = 39; Figure 2B), indicating that both ring closure velocity and time lag control asymmetric ring closure in normal cells.

**Figure 2.**
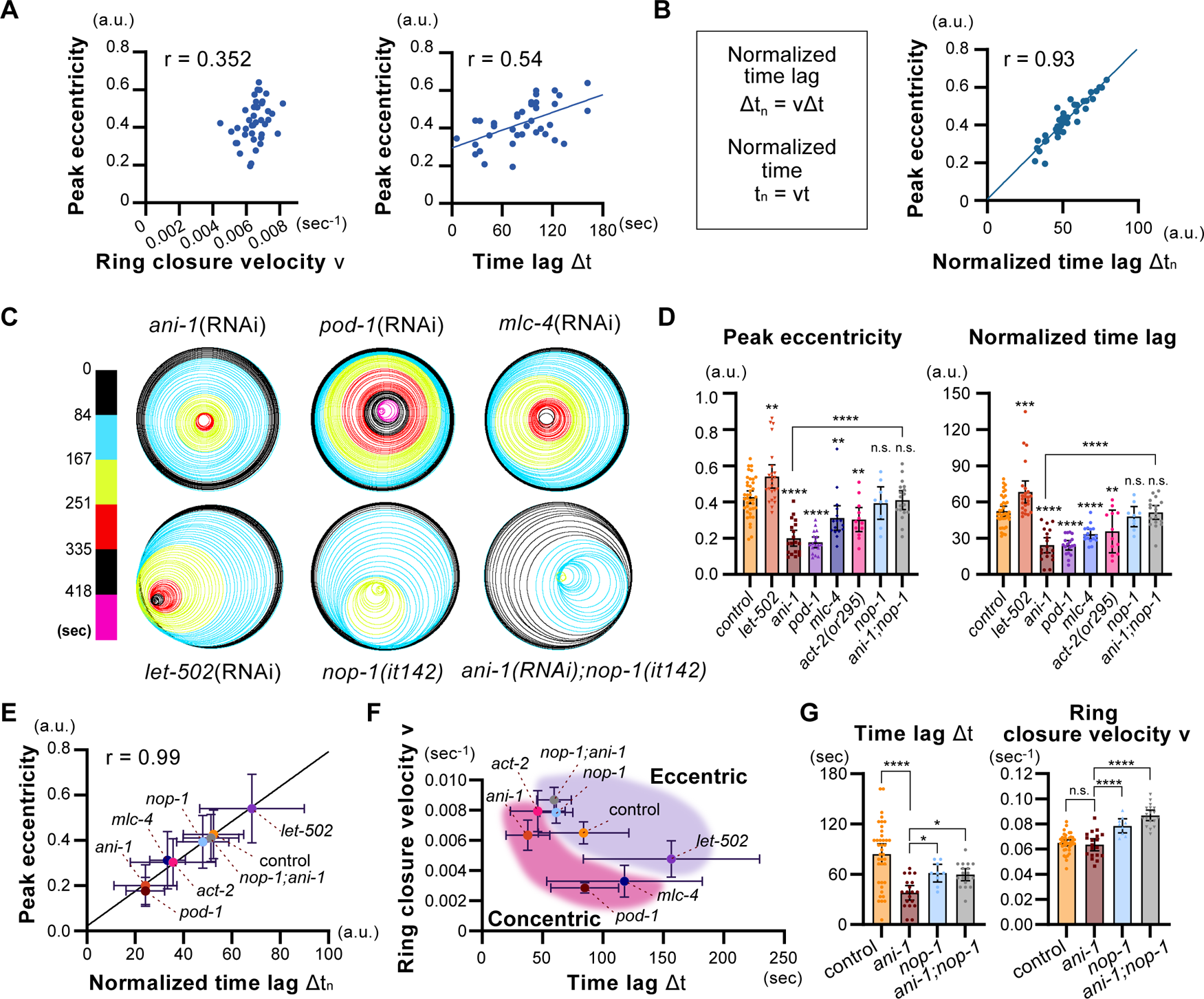
Ring closure velocity and a delay in lagging edge furrowing collectively determine the ring eccentricity. (**A**) Relationship between peak eccentricity, ring closure velocity, and time lag in normal cells. (**B**) Relationship between peak eccentricity and normalized time lag in normal cells. (**C**) Ring trajectories of different RNAi and mutant backgrounds. (**D**) Peak eccentricity and normalized time lag in different backgrounds. (**E**) Relationship between peak eccentricity and normalized time lag in different backgrounds. (**F**) The relationship between ring closure velocity and time lag in different backgrounds. (**G**) Time lag and ring closure velocity. Times are relative to cytokinesis onset. Error bars indicate 95% confidence intervals except for panel E and F, in which standard deviations were shown. The value r represents Pearson’s correlation coefficient. *P*-values were calculated using one-way ANOVA followed by Holm–Sidak’s multiple comparison test. Scale bars, 10 µm.

To investigate whether ring closure velocity and time lag generally control eccentricity, we analyzed these parameters in mutants or RNAi knockdowns of actomyosin regulators (Figures 2C–G and S2A–F). In addition to the previously reported ANI-1/Anillin, we found that POD-1/Coronin and MLC-4/myosin regulatory light chain are required for ring eccentricity (Figure 2C–D). We also confirmed that a deletion mutation of the worm-specific RhoA activator NOP-1 rescued the *ani-1(RNAi)* phenotype ^35^ and that the knockdown of LET-502/ROCK did not cause eccentricity defects ^6^, as reported previously (Figure 2C–D).

We calculated time lag, ring closure velocity, and normalized time lag in these conditions. Time lag (raw values) and peak eccentricity were not particularly correlated (Pearson’s r = 0.54, *p* = 0.21; Figure S2G). However, we found that normalized time lag is strongly correlated with peak eccentricity (Pearson’s r = 0.99, p < 0.0001; Figure 2E). The relationship between ring closure velocity and time lag also shows that conditions with asymmetric closure and symmetric closures are separated within the parameter space (Figure 2F). Therefore, cells that exhibit faster ring closure and longer time lag tend to display asymmetric ring closure. Using these data, we investigated why the *nop-1* mutation was able to rescue the *ani-1(RNAi)* phenotype. We found that the *nop-1* mutation increased both time lag and ring closure velocity compared to *ani-1(RNAi)* alone (Figure 2G), suggesting that the rescue is due to the increase of these parameters. These results suggest that ring eccentricity is collectively determined by the ring closure velocity and a delay in furrowing at the lagging edge.

### Reduced ring-directed cortical flow at the lagging cell cortex accompanies circumferential-axis cortical flow

We analyzed the mechanism of the lagging edge furrowing delay. As ring-directed cortical flow is proposed to contribute to the positive feedback regulation of the contractile ring assembly (Figure 3A)^37^, we hypothesized that the delay is due to the feedback loop having lower activity at the lagging edge than at the leading edge. To test this hypothesis, we analyzed cortical flow at the leading and lagging cortices using particle image velocimetry (PIV) ^43^ (Figures 3B and S3, and Movie S2).

**Figure 3.**
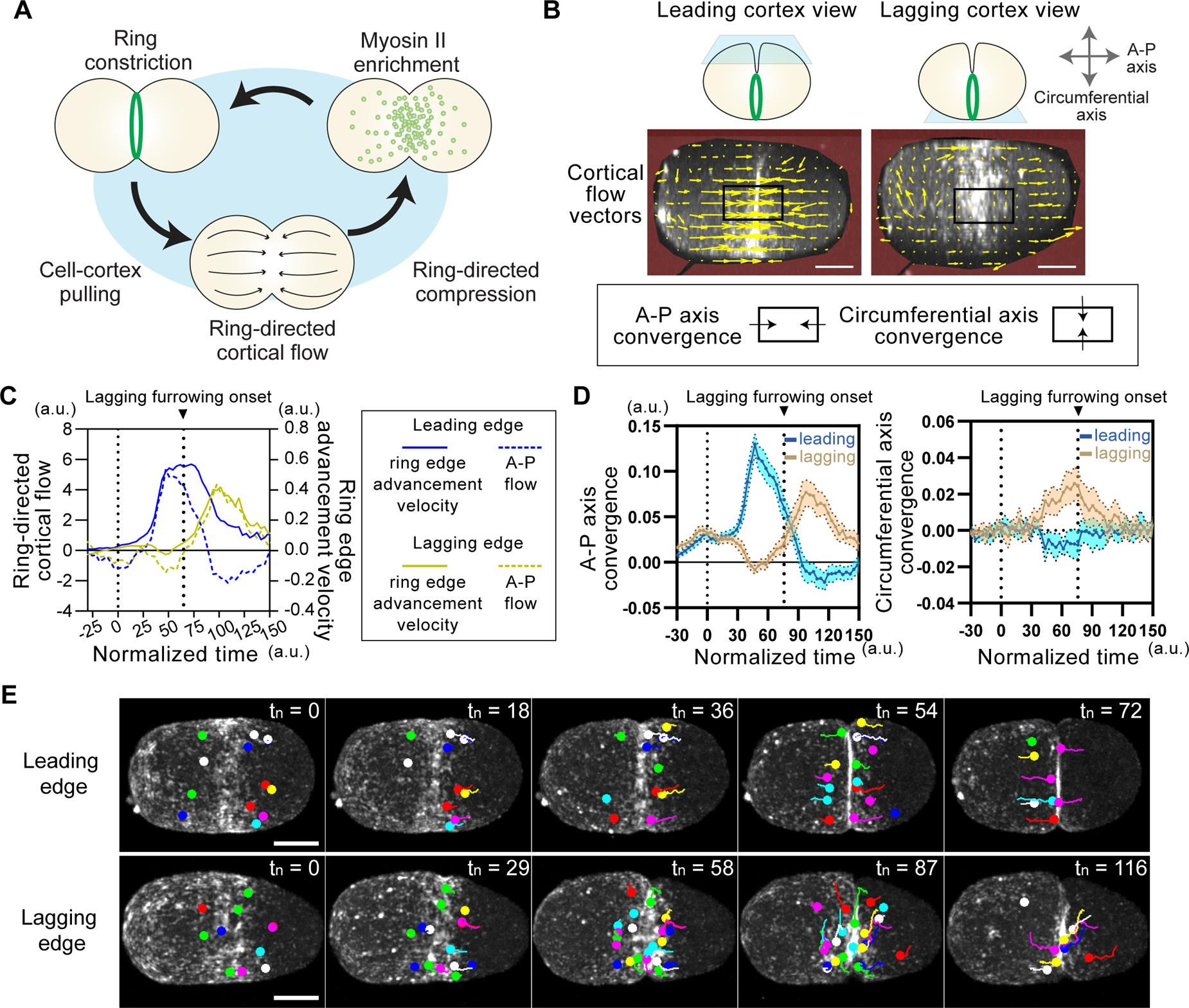
Reduced ring-directed cortical flow at the lagging cell cortex accompanies circumferential-axis cortical flow. (**A**) A cortical flow-dependent positive feedback model in the contractile ring assembly. Ring-dependent cell cortex pulling generates ring-directed cortical flow and induces ring-directed compression. Resulting myosin enrichment to the ring further facilitates ring constriction. (**B**) Measurement of cortical flow. Leading and lagging cortex views were generated by computationally rotating 4D timelapse data. Cortical flow vectors were estimated using particle image velocimetry (yellow arrows). Convergence of cortical flow vectors were calculated to estimate the influx of cortical flow within the equatorial ROI (black boxes). (**C**) The relationship between ring edge advancement and ring-directed cortical flow. Cortical flow in the area anterior to the ring is shown. (**D**) A-P axis convergence and circumferential axis convergence in leading and lagging cell cortices. (**E**) Myosin foci tracking during cytokinesis. Scale bars, 10 µm.

First, we analyzed the mechanism of ring-directed cortical flow generation, which is an unclear part of the proposed feedback model. Cortical flow is a phenomenon driven by a gradient in actomyosin contractility ^44^. This gradient is established during cytokinesis, with higher contractility at the equator and relaxation at the pole ^23, 37, 38^. Given that the contractility gradient continues to exist during ring assembly and closure, the ring should consistently pull the cell cortex from the polar region. To confirm this prediction, we plotted velocities of ring edge advancement (Figure 3C; solid lines) and ring-directed cortical flow (Figure 3C; dotted lines) and found that they correlate well both at the leading and lagging edges (leading edge: Pearson’s r = 0.77, *p* < 0.0001, lagging edge: Pearson’s r = 0.89, *p* < 0.0001). Furthermore, ring closure velocity and peak ring-directed flow are highly correlated in different RNAi and mutant conditions, including *ani-1(RNAi)* (Figure 8D; Pearson’s r = 0.94, *p* = 0.002). These results suggest that the invaginating ring edge consistently pulls the nearby cell cortex to generate ring-directed cortical flow to form a positive feedback loop (Figure 3A).

Next, we analyzed the difference in cortical flow between leading and lagging cortices. To better understand the influx of the cortical flow within the equatorial region, we calculated the convergence of cortical flow vectors along the A-P axis and the orthogonal, circumferential-axis (Figure 3B). Furthermore, we tracked myosin foci to confirm the PIV data (Figure 3E). At the leading cell cortex, A-P axis convergence increased rapidly and reached to its peak before the onset of lagging edge furrowing (Figure 3D; left). Consistently, myosin foci tracking showed continuous ring-directed cortical flow (Figure 3D; top). Conversely, A-P axis convergence in the lagging cell cortex decreased shortly after cytokinesis onset and reached a local minimum before the onset of lagging edge furrowing (Figure 3D; left, Figure 3E; t_n_ = 58). During this period, we observed increase in circumferential axis convergence only at the lagging cell cortex (Figure 3D; right). These results suggest that circumferential-axis cortical flow negatively correlates with ring-directed cortical flow.

### Cells undergoing asymmetric ring closure exhibit polarized mechanical loading at the lagging cortex

Circumferential cortical flow could inhibit ring-directed cortical flow, potentially through a mechanical process (Figure 3A). To infer cortical mechanics from the flow vectors, we used cortical convergence. Before furrowing onset, convergence at the surface signifies cortical compression (Figure 4A, top). After the onset of furrow ingression, convergence at the surface represents the sum of cortical compression and ingression, similar to how converging surface sea water sinks towards the ocean floor (Figure 4B, bottom).

**Figure 4.**
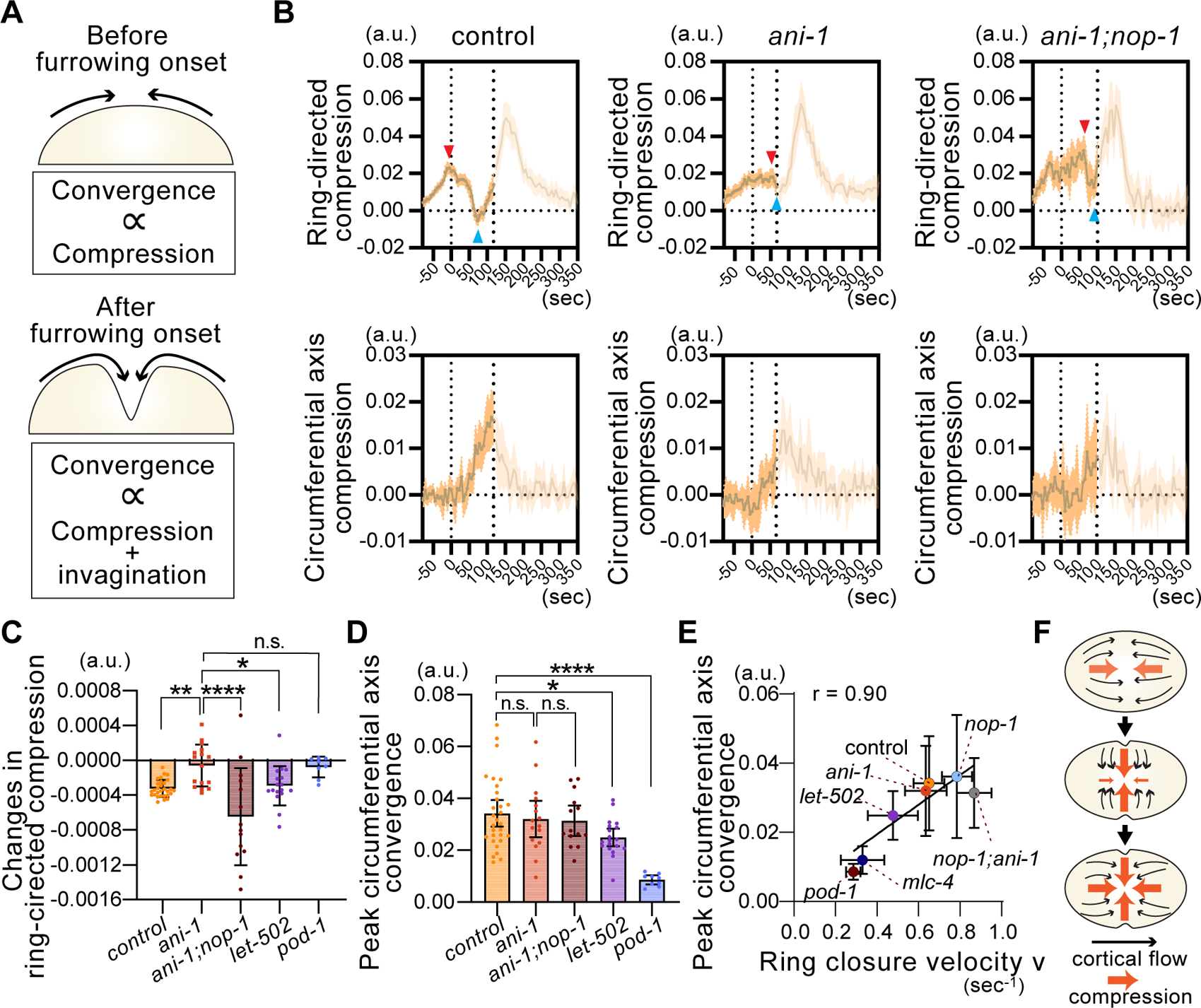
Cells undergoing asymmetric ring closure exhibit polarized mechanical loading at the lagging cortex. (**A**) Inferring cortical mechanics from cortical convergence data. Before furrowing onset, cortical convergence is proportional to the cortical compression. After furrowing onset, convergence is proportional to the sum of compression and invagination. (**B**) Cortical compression at the lagging equatorial cell cortex. Mean cortical flow convergence before the onset of lagging edge furrowing is shown. The transparent parts of graphs show the convergence after the furrowing onset. Red and blue arrowheads indicate local maxima and minima between cytokinesis onset (left dotted line) and lagging edge furrowing onset (right dotted line) in each graph. (**C**) Reduction rate of ring-directed compression at the lagging equatorial cortex. (**D**) Peak circumferential axis convergence observed at the lagging equatorial cortex. (**E**) Relationship between peak circumferential axis convergence and ring closure velocity. The value r represents Pearson’s correlation coefficient. (**F**) Summary of the cortical flow and cortical compression at the lagging equatorial cell cortex. Ring-directed compression decreases as the circumferential compression increases. P-values were calculated using one-way ANOVA followed by Holm-Sidak’s multiple comparison test. Times are relative to cytokinesis onset. Error bands and bars indicate 95% confidence intervals. Scale bars, 10 µm.

In control embryos, we observed that ring-directed cortical compression at the lagging cell cortex decreased upon cytokinesis onset (Figures 4B; top left). However, this decline was not apparent in *ani-1(RNAi)* (Figure 4B; top middle). The decline reemerged with the introduction of a *nop-1* mutation (Figure 4B; top right). On the other hand, there were no marked differences in circumferential axis convergence under these conditions (Figure 4B; bottom row). The rate of reduction in ring-directed cortical compression, estimated from the lines connecting local maxima and minima (red and blue arrowheads in the Figure 4B), also confirmed a similar relationship (Figure 4C). We found that conditions with asymmetric ring closure (control, *ani-1;nop-1*, and *let-502*) and symmetric ring closure (*ani-1* and *nop-1*) exhibited a reduction and no change in ring-directed compression, respectively (Figure 4C). Since control and *ani-1(RNAi)* displayed a similar ring closure velocity (Figure 2F), the observed cortical relaxation cannot be attributed to the ring closure rate. In contrast, peak circumferential-axis convergence strongly correlates with ring closure velocity (Pearson’s r = 0.90, *p* = 0.005) (Figure 4D–E). These results suggest that in general, contractile ring closure induces circumferential axis compression. This compression is then accompanied by reduced ring-directed compression in cells undergoing asymmetric ring closure (Figure 4F). However, ring-directed compression in *ani-1(RNAi)* does not respond to the compression along the circumferential axis.

### Non-ring-directed cortical compression suppresses ring-directed cortical flow

We hypothesized that any non-ring-directed cortical compression, like the circumferential-axis compression we observed, inhibits ring-directed cortical flow. To test this hypothesis, we induced ectopic cortical compression using a semi-dominant temperature-sensitive mutation of actin, *act-2(or295)*, known to cause actin filament stabilization and ectopic cortical contractility ^45^ (Figure 5A). In normal cells, the leading cortex demonstrates consistent ring-directed flow, while the lagging cortex experiences a period of delay followed by ring-directed flow (Figure 5A–B). In *act-2(or295)*, ring-directed movement of myosin foci was suppressed during contraction and pronounced during relaxation (Figure 5C). Ring closure is necessary for relaxation-induced ring-directed cortical flow, as similar flow was not observed post-cytokinesis (Figure 5D). These results suggest that non-ring-directed cortical compression inhibits ring-directed cortical flow at the lagging cell cortex (Figure 5E).

**Figure 5.**
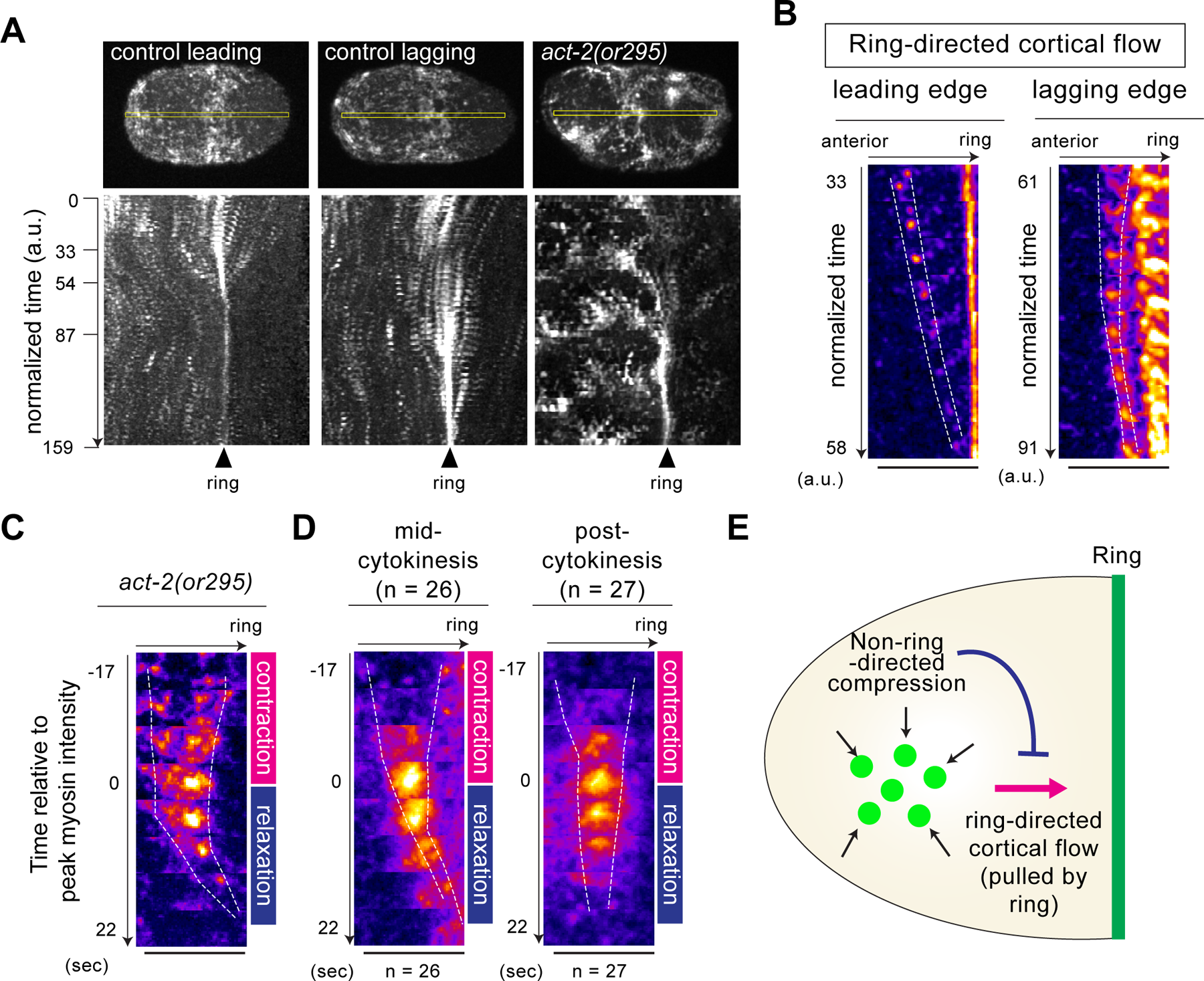
Non-ring-directed cortical compression suppresses ring-directed cortical flow. (**A**) Kymographs of cortical myosin. Yellow boxes in the top images were used to generate kymographs. (**B**) Ring-directed cortical flow in control zygotes. Leading edge shows consistent flow while the lagging edge flow is preceded by the stabilization. (**C**) Ring-directed cortical flow in *act-2* mutant zygotes. A myosin focus movement during contraction and relaxation of the area behind the ring was shown. (**D**) Averaged ring-directed flow in *act-2* mutant zygotes. (**E**) The mechanism of compression-dependent suppression of the ring-directed cortical flow. While ring-directed compression promotes ring assembly, non-ring-directed compression suppresses ring-directed cortical flow. Times are relative to cytokinesis onset. Scale bars, 10 µm.

### Non-ring-directed compression delays ring edge advancement without suppressing ring constriction

A previous study demonstrated that laser-induced ablation of the cell cortex behind the ring did not accelerate the ring closure rate ^37^. This suggests that ring closure is not constrained by the mechanical resistance of the cell cortex behind the ring. If this is indeed the case, the observed retardation of ring-directed flow would not limit the ring closure rate, and therefore, would not influence the ring eccentricity. We tested this possibility using highly contractile *act-2(or295)* mutants and found that the overall ring closure curves were similar to those of the control, with the ring velocity even exceeding than the control (Figure 2F). This confirms the first part of our prediction. However, the contractile rings of the *act-2* mutant frequently displayed oscillatory movement during constriction and exhibited a lower mean peak eccentricity compared to the control (Figures 6A and 2D). These results suggest that cortical compression influences ring position without altering the constriction rate.

**Figure 6.**
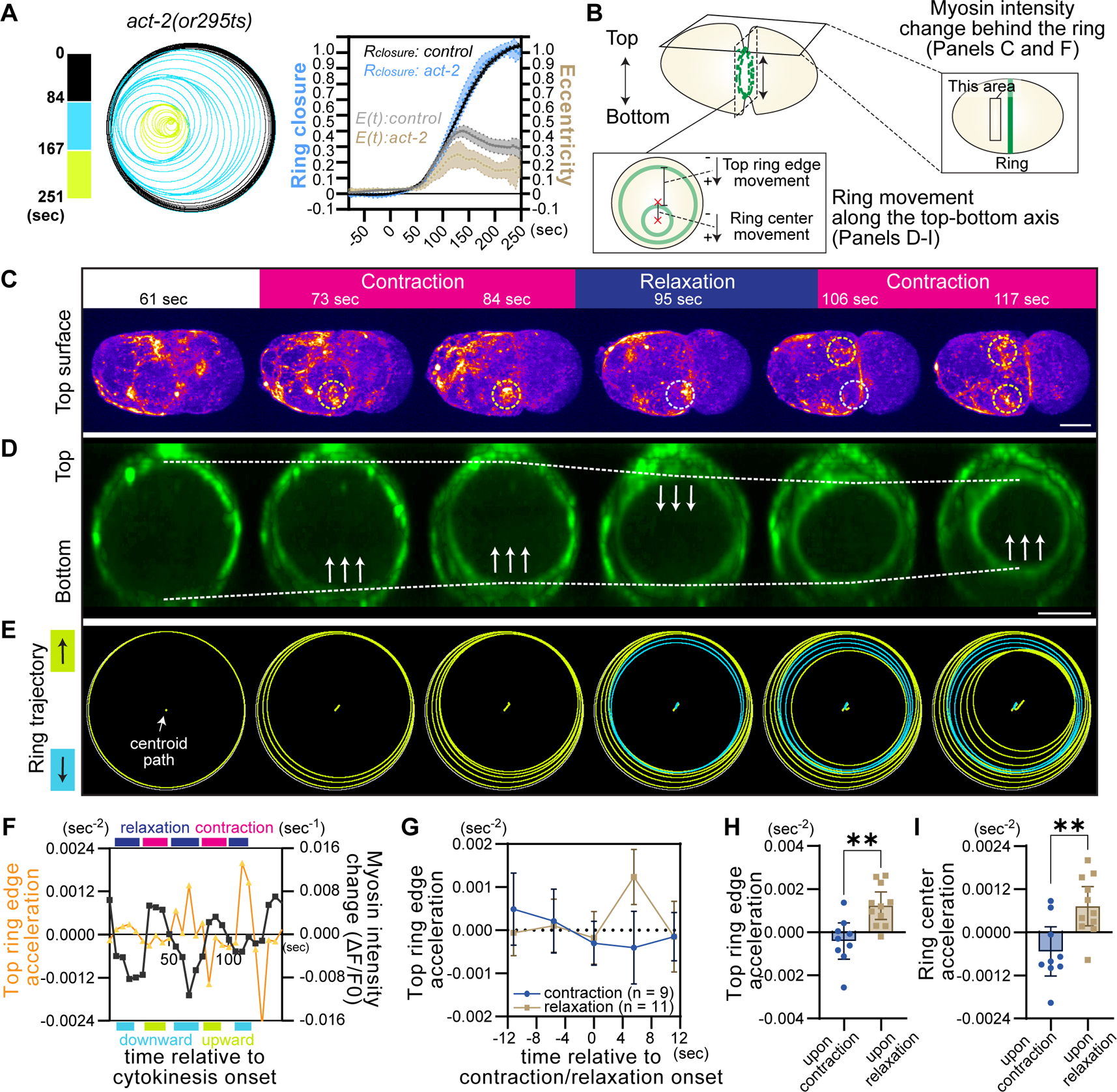
Non-ring-directed cortical compression delays ring edge advancement without suppressing ring constriction. (**A**) Contractile ring trajectory of a single *act-2(or295ts)* mutant (left panel) and the mean values of ring closure and eccentricity (graph). (**B**) Planes shown in (C–E). Changes in myosin intensity indicate contraction and relaxation. (**C**) NMY-2::GFP at the top surface during cytokinesis. Yellow and light blue dotted circles indicate the area exhibiting contraction and relaxation, respectively. (**D**) Ring en face view of the same embryo shown in panel C. Dotted lines indicate top and bottom limits of the contractile ring. Arrows indicate the movement of the ring limit. (**E**) Ring trajectories of the contractile ring shown in (D). Top- and bottom-oriented movements are shown in yellow and blue, respectively. (**F**) Relationship between top ring acceleration and myosin intensity changes. Positive ring edge acceleration represents ring ingression toward the bottom surface. (**G**) Top ring edge acceleration upon contraction and relaxation. (**H**) Top ring edge acceleration measured 5.6 s post-contraction and relaxation. (**I**) Ring center acceleration 5.6 s after contraction and relaxation. Error bars and bands indicate 95% confidence intervals. Scale bars, 10 µm.

We investigate how cortical compression behind the ring affects ring positioning. To infer cortical compression, we measured myosin intensity changes at the cell surface closest to the objective lens (referred to as the top surface) (Figure 6B). The upward and downward movements of contractile ring along the top-bottom axis were also measured using the same samples (Figure 6B). During contraction at the top surface, the downward movement of the top ring edge was often halted, as though the top ring edge were anchored (Figure 6C–E; labeled as “contraction,” Movie S3). Conversely, relaxation of the top surface often coincided with the bottomward movement of the top ring edge (Figure 6C–E; labeled as “relaxation,” Movie S3). Quantitative analysis of cortical compression and acceleration of the top ring edge also confirmed these trends (Figure 6F–H; positive acceleration indicates movement towards the bottom surface). These findings suggest that cortical compression behind the ring impedes the advancement of the nearby ring edge.

How does the compression-dependent retardation of the ring edge occur without delaying the overall ring constriction rate? Notably, we frequently observed upward movement of the bottom ring edge when the top surface is compressed (Figure 6C–E, I). Thus, it is likely that when a certain part of the ring edge advancement is prevented, other areas of the ring edge pull more cortex from the nearby surface, maintaining a similar constriction rate.

### Artificial inhibition of ring-directed cortical flow is sufficient to induce ring eccentricity

Our data suggest that the retardation of ring-directed cortical flow can trigger asymmetric ring closure. To directly test this hypothesis, we inhibited ring-directed cortical flow in symmetrically dividing cells. It is known that the two-cell stage AB cell undergoes asymmetric ring closure, similar to the P_0_ zygote ^46^ (Figures 7A and S5). We found that unilateral ring closure requires cell contact with the neighboring P_1_ cell (Figure 7B–C). To inhibit ring-directed cortical flow, we attached adhesive polystyrene beads to the symmetrically dividing isolated AB cell (Figure 7D–E). The beads, which were coated with positively charged Rhodamine fluorescent dyes and salts, adhere to the plasma membrane. This adhesion is presumed to occur through electrostatic interaction, similar to the mechanism observed with poly-L-lysine coating. As shown in our previous study, attachment of the 30 µm diameter beads reduced ring-directed flow in the area proximal to the bead attachment site (Figure 7E) ^1^. The response scaled with the bead diameter, suggesting that passive mechanical resistance imposed by adhesive beads limits ring-directed flow (Figure 7E). Notably, the attachment of the 30 µm beads increased peak eccentricity, and about 50% of the cells exhibited a level of ring eccentricity comparable to that of the intact AB cell *in vivo* (Figure 7C, F–H). Thus, we conclude that mechanical retardation of ring-directed cortical flow is sufficient to induce asymmetric ring closure.

**Figure 7.**
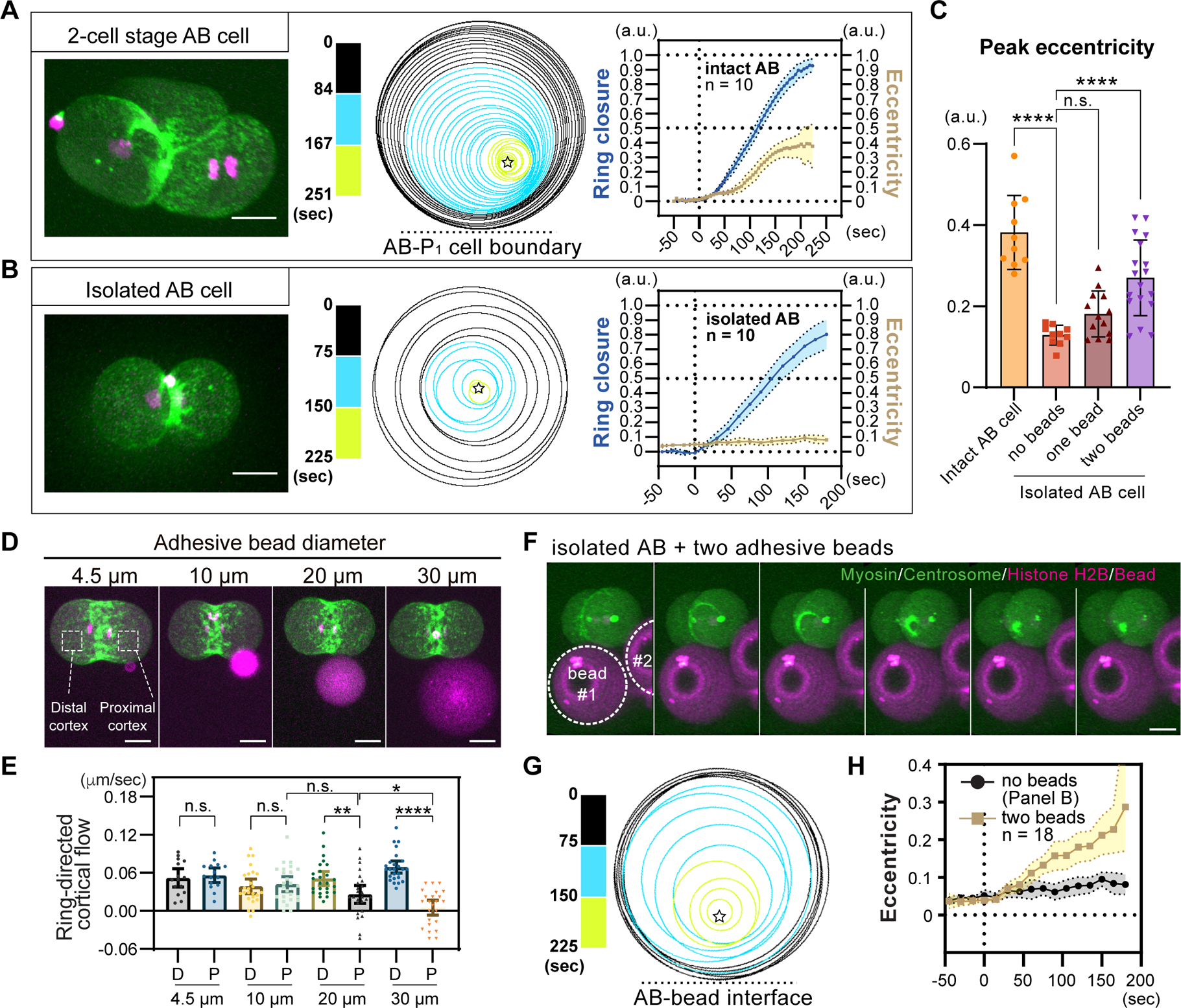
Artificial inhibition of ring-directed cortical flow is sufficient to induce asymmetric ring closure. (**A**, **B**) Contractile ring closure in the intact and isolated AB cell. NMY-2::GFP and mCherry::histone H2B (left), ring trajectories (middle), and ring closure and eccentricity curves (right) are shown. (**C**) Mean peak eccentricity of the contractile ring in the AB cells. (**D**, **E**) Attachment of different sized adhesive beads to the isolated AB cell. Ring-directed cortical flow were plotted for proximal and distal to the bead attachment site in E. (**F**–**H**) Oblique view of contractile rings in the bead attached AB cell. Ring trajectory and mean eccentricity are shown in G and H, respectively. Times are relative to cytokinesis onset. Error bands and bars indicate 95% confidence intervals. Scale bars, 10 µm.

### Anillin tunes contractile ring mechanosensitivity

Our analysis suggests that contractile rings in both the P_0_ zygote and the two-cell stage AB cell exhibit mechanosensitivity. We found that *ani-1(RNAi)* embryos exhibited reduced ring eccentricity in both cell types, whereas *pod-1(RNAi)* showed defects only in P_0_ (Figure 8A; top panel). The observed loss of asymmetric closure in *pod-1(RNAi)* can be explained by reduced circumferential-axis convergence (Figure 4D and 8A). These results suggest that the contractile ring in *pod-1(RNAi)* retains its mechanosensitivity but lacks the P_0_ mechanical cue. Conversely, *ani-1(RNAi)* embryos had a normal level of circumferential-axis convergence (Figure 4D and 8A), indicating that the contractile rings in *ani-1(RNAi)* have impaired mechanosensitivity.

**Figure 8.**
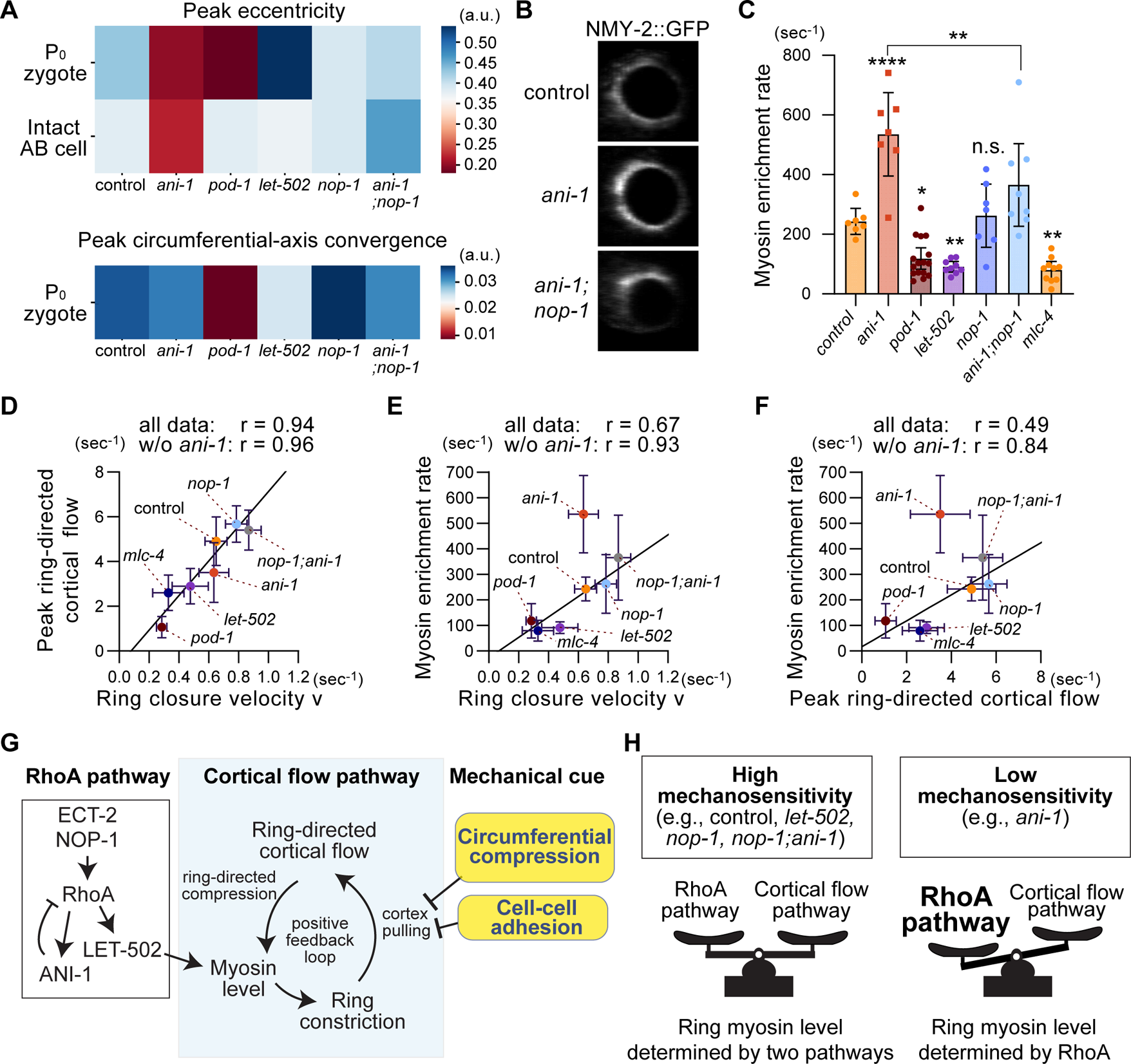
Anillin tunes contractile ring mechanosensitivity. (**A**) Heatmaps of peak eccentricity and peak circumferential-axis convergence. *ani-1(RNAi)* embryos exhibit defects in asymmetric ring closure in both P_0_ and AB cells. (**B**, **C**) Contractile ring myosin enrichment. Total myosin signal intensity in the ring (B) was used to estimate myosin enrichment rate in (C). (**D**-**F**) Relationship among ring closure velocity, peak ring-directed cortical flow, and myosin enrichment rate. (**G**) Proposed model of contractile ring-dependent mechanosensation. (**H**) Proposed model of Anillin-dependent tuning of contractile ring mechanosensitivity. Error bars indicate 95% confidence intervals. Scale bars, 10 µm. p-values were calculated by one-way ANOVA with Holm-Sidak’s multiple comparison test.

We observed that contractile rings in *ani-1(RNAi)* often retain a unusually high level of myosin signal (Figure 8B). Indeed, the myosin enrichment rate in *ani-1(RNAi)* is greater than any of the other RNAi groups that we tested (Figure 8C). Therefore, we reasoned that the ectopic enrichment of myosin might disrupt the mechanosensitivity of the *ani-1(RNAi)* ring. We hypothesized that a cortical flow-dependent positive feedback loop in the contractile ring assembly sensitizes the contractile ring to mechanical stresses, and that the feedback loop is disrupted in *ani-1(RNAi)* embryos (Figure 8G; central box).

We observed a strong correlation between ring closure velocity and peak ring-directed cortical flow across different RNAi/mutant groups including *ani-1(RNAi)*, as described above (Figure 8D). We also found that the myosin enrichment rate is highly correlated with ring closure velocity and peak ring-directed cortical flow among sample groups excluding *ani-1(RNAi)* (Figure 8E: Pearson’s r = 0.93, *p* < 0.007, Figure 8F: Pearson’s r = 0.84, p < 0.03). Conversely, *ani-1(RNAi)* exhibited excess myosin enrichment rate relative to the ring closure velocity and peak ring-directed cortical flow (Figure 8E–F). These results suggest that the positive feedback loop between ring-directed cortical flow, myosin enrichment, and ring constriction was disrupted in *ani-1(RNAi)* embryos.

We further observed that the *nop-1* mutation rescued the asymmetric ring closure defects in *ani-1(RNAi)* in both the P_0_ and AB cells (Figure 8A), indicating that the *nop-1* mutation restored the impaired mechanosensitivity of the contractile ring. The rescue of polarized mechanical loading also supports this idea (Figure 4B–C). Analysis of myosin levels revealed that the *nop-1* mutation reduced the ectopic myosin enrichment rate in *ani-1(RNAi)* (Figure 8B–C). As a result, *ani-1(RNAi); nop-1(it142)* embryos regained a normal level of myosin enrichment relative to ring closure velocity and peak ring-directed flow (Figure 8D–F). Therefore, we concluded that impairment of the contractile ring’s mechanosensitivity is due to the disruption of the positive feedback loop caused by excessive myosin enrichment.

## Discussion

In this study, we have identified the mechanisms underpinning contractile ring mechanosensation and the tuning of mechanosensitivity. Previous studies have shown that ring-directed cortical compression promotes ring assembly, but our findings demonstrate that non-ring-directed compression slows down ring-directed cortical flow and the advancement of the ring edge. The suppression of ring-directed cortical flow was sufficient to induce asymmetric ring closure in symmetrically dividing cells. The mechanism of ring mechanosensation relies on a normal rate of myosin enrichment, in relation to the velocities of ring closure and cortical flow. Finally, our results show that anillin is required for the sensitization of the contractile ring against intercellular and intracellular mechanics.

Based on our analysis and previous studies, we propose a cortical flow-dependent contractile ring mechanosensation model (Figure 8G). The critical component of the model is the positive feedback loop among ring constriction, ring-directed cortical flow, and myosin enrichment. Khaliulin et al. (2018) proposed this feedback loop in the contractile ring constriction process based on correlations among these parameters in normal zygotes ^37^. Our study further confirms these correlations in RNAi and mutant backgrounds with varying ring closure velocities (Figure 8D–F). In our model, intracellular and intercellular mechanical cues locally limit ring-directed cell cortex-pulling at the lagging cell cortex (Figure 8G; arrows from the yellow boxes), thereby suppressing the ring-directed cortical flow. The local suppression of the ring-directed cortical flow should reduce compression-dependent myosin enrichment, resulting in the formation of the structurally asymmetric ring observed in previous studies ^6^. The local reduction in myosin levels should then locally slow down ring edge advancement, resulting in asymmetric ring closure.

Our analysis underscores the central role of the cortical flow pathway in ring mechanosensation. Conversely, the RhoA pathway likely inhibits the mechanosensation process. We found that reduced activities of the RhoA activator NOP-1 and effector LET-502 did not influence ring eccentricity. However, we showed impaired mechanosensitivity in *ani-1(RNAi)* embryos, in which myosin was highly enriched in the contractile ring (Figure 8). Recent research has shown that ANI-1-binding proteins GCK-1 and CCM-3 negatively regulate RhoA activity by promoting RhoGAP cortical localization ^47^. Therefore, it is likely that ANI-1 knockdown lifted this negative regulation, leading to the activation of RhoA-dependent myosin enrichment. Indeed, normal myosin enrichment as well as ring mechanosensitivity were restored in *ani-1; nop-1* (Figure 8A–C). We propose that the balance between RhoA-dependent and cortical flow-dependent myosin enrichment tunes ring mechanosensitivity (Figure 8H). The mechanosensitivity of the ring should decrease when the contribution of the cortical flow pathway-dependent myosin enrichment is negligible compared to that of RhoA-dependent pathway (Figure 8H).

The structural asymmetry of the contractile ring in the *C. elegans* zygote likely arises from asymmetric activity in the cortical flow-dependent feedback mechanisms between the leading and lagging cell cortices ^37^. A similar structural asymmetry has been reported in *Drosophila* pupal dorsal thorax epithelia ^11^. However, *Drosophila* embryonic epithelial cells ^2^, follicular epithelia ^34^, and pupal wing epithelia ^11^ do not display such structural asymmetry. As our study identified that the cortical flow-dependent feedback mechanism mediates ring mechanosensation, this variability suggests that the mechanosensitivity of contractile ring can be tuned differently during development, enabling tissue-specific morphogenic cytokinesis. The cortical flow pathway likely translates subtle differences in tissue mechanics into asymmetry in the furrowing rate, while excessive RhoA activity could suppress ring-dependent mechanosensing. Cell types that lack structurally asymmetric rings may undergo asymmetry ring closure by relying on extreme differences in tissue stiffness (e.g., a stiffer apical cortex connected to adherens junction vs a softer basal cortex) rather than on high ring mechanosensitivty. Alternatively, these cell types might employ other mechanosensitive mechanisms, such as mechanosensitive recruitment of myosin regulators to neighboring cells ^48, 49^.

The tuning of ring mechanosensitivity would be critical in animal development. Here, we will use lumen morphogenesis as an example to explain this concept. A previous study showed that 3D cultures of epithelial cells initially undergo symmetric cytokinesis to form a central lumen ^12, 50–53^. Subsequent ring closures are asymmetric, maintaining a solitary lumen. In the case of intraflagellar transport protein IFT88 knockdown, the first cytokinesis becomes asymmetric, leading to the formation of multiple lumens ^12^. Consistently, a mouse model with a loss of function mutation of *ift88* exhibits a polycystic kidney phenotype ^54^. In this context, the contractile ring in the first cytokinesis should exhibit lower mechanosensitivity to prevent ectopic unilateral cytokinesis, while the contractile ring in subsequent cytokinesis should be highly mechanosensitive. Our study provides a theoretical framework for a deeper understanding of tissue-specific mechanochemical regulation of the contractile ring, which is crucial for further elucidating the role of cytokinesis in these morphogenetic events.

## Acknowledgements

We thank Kota Mizumoto and Don Moerman for critical reading of the manuscript. Some strains used in this study were obtained from the *Caenorhabditis* Genetics Center (funded by the NIH Office of Research Infrastructure Programs; P40 OD010440). We also thank Bruce Bowerman for providing valuable advice, Chris Doe for sharing lab equipment, and the Sugioka lab members for general discussions. This work was supported by the Canadian Institutes of Health Research (Project Grant; PJT-169145), Government of Canada’s New Frontiers in Research Fund (NFRFE-2019-00310), and the Health Research BC (Scholar Award; SCH-2020-0406) to K.S. C.R.H was supported by British Columbia Graduate Scholarship.

## Author contributions

**C.R.H.**: Formal analysis, Investigation, Writing-Original Draft, Validation, Visualization. **G.S.**: Formal analysis, Investigation, Validation, Visualization. **W.F.**: Formal analysis, Investigation. **J.Z.**: Formal analysis, Investigation. **K.S.**: Conceptualization, Formal analysis, Investigation, Methodology, Software, Validation, Writing-Original Draft, Writing-Review & Editing, Supervision, Funding acquisition.

## Methods

### Experimental Model and Subject Details

All *C. elegans* strains were cultured using the standard method ^55^. A temperature sensitive-actin mutant *act-2(or295)* was cultured at 15 °C until the L4 larval stage and incubated at 25 °C overnight before imaging. The following transgenes were used: *cp13*[*nmy-2*::GFP + LoxP] (non-muscle myosin II) ^56^, *or1940*[GFP::*sas-7*] (centriole marker) ^57^, *itIs37* (mCherry::histone H2B), *ruIs32*[*pie-*1p::GFP::H2B::*pie-1* 3’UTR + *unc-119*(+)].

### RNAi

Feeding RNAi was performed at 25 °C using the standard method ^58^. For control RNAi, a bacterial strain carrying an empty vector (L4440) was used. To observe *let-502(RNAi)* P_0_ zygotes, the L2 stage larvae were cultured on freshly prepared feeding RNAi plates on day 1. The L4 larvae were then transferred to new feeding RNAi plates on day 2 and imaged on day 3. For observing *let-502(RNAi)* AB cell, we performed L4 RNAi to avoid abnormal AB spindle orientation. For *pod-1* and *mlc-4*, L4 larvae were cultured on feeding RNAi plates and used for imaging on the next day.

### Blastomere isolation

Blastomeres were isolated as described before ^59^, with some modification ^60^. We cut the gravid adult worms in egg salt buffer and treated them with hypochlorite solution [75% Clorox (Clorox) and 2.5N KOH] for 50 s. After washing twice with Shelton’s growth medium ^61^, embryos were transferred to fresh Shelton’s growth medium. Eggshell and permeability barrier were removed by mouth pipetting with hand-drawn microcapillary tubes (10 μL, Kimble Glass Inc.). The two-cell stage eggshell-free embryos were further pipetted to remove the cell-cell contact.

### Adhesive polystyrene bead preparation

The detailed method is described in our previous papers ^1, 60^. Approximately 10 mg carboxyl-modified polystyrene beads with diameters of 30 µm (Kisker Biotech GmbH & Co.), 20 µm, 10 µm, and 4.5 µm (Polysciences) were washed twice with 100 mM 2-(N-morpholino) ethanesulfonic acid (MES) buffer (pH6.5) and incubated with 1 mL MES buffer containing 10 mg 1-Ethyl-3-(3-dimethylaminopropyl) carbodiimide (EDAC) for 15 min at room temperature (22.5°C). We washed the beads twice with phosphate-buffered saline (PBS) and incubated them with 0.5 mL PBS containing 0.05 µg Rhodamine Red-X succinimidyl ester (ThermoFisher Scientific) for 5 min. The appropriate concentration was also determined by treating the beads with a series of serially diluted Rhodamine Red-X solution ^60^. The beads were washed twice with PBS and stored in PBS at 4 °C. Adhesiveness of the beads was confirmed by attaching them to the isolated blastomere using the mouth pipette. If successful, the adhesion is firmly established and cells do not dissociate spontaneously.

### Live-imaging sample preparation

To obtain embryos, gravid adults were dissected on a coverslip, in a droplet of 10–12 µl of refractive index-matching medium (30% iodixanol diluted in egg salt buffer, supplemented with 30 µm diameter plain polystyrene beads) as described before ^42^. After placing the coverslip gently onto a slide glass, three edges of coverslip were sealed with petroleum jelly (Vaseline), with one edge remaining open to the air. We realized that this method improved the success rate of imaging for inexperienced users. Inexperienced users may observe cell death due to the acute compression during the sample preparation and require training using control strains. If all the processes were performed correctly, the imaging condition does not have adverse effects on the embryonic viability, as judged by normal cell division in the next cell cycle.

### Microscopy

Intact embryos were imaged using a microscope Olympus IX83 (Olympus), equipped with a spinning-disk confocal unit CSU-W1 (Yokogawa), a scientific CMOS camera Prime 95B (Photometrics), a piezoelectric stage NANO-Z (Mad City Labs), a silicon immersion objective UPLSAPO60XS2 (NA1.3, 60X; Olympus), and a beam splitter Optosplit II (Cairn Research), which is controlled by Cellsense Dimension (Olympus). A silicone immersion oil (Z81114; refractive index: 1.406 at 23 °C; Olympus) was used as an immersion medium. Samples were illuminated by diode-pumped lasers with 488 nm and 561 nm wavelengths, and the simultaneous two color-imaging was performed with 150 ms camera exposure time, 1 µm Z-step size with a total of 31 slices per frame, 5.6-s interval, and the duration of 15 min. Isolated blastomeres were imaged using a microscope Leica DMi8 (Leica Microsystems), equipped with a spinning-disk confocal unit CSU-W1 with Borealis (Andor Technology), dual EMCCD cameras iXon Ultra 897 (Andor Technology), and an oil-immersion PL APO objective lens (NA1.4, 63X; Leica), and controlled by Metamorph (Molecular Devices). Data in Figure 7B and 7F were imaged with 1.5 µm Z-step size and 15-s intervals. Data in Figure 7D were imaged with 1.5 µm Z-step size and 10-s intervals, with only the half volume closer to the objective lens imaged.

### Quantification of the P_0_ contractile ring dynamics

The obtained 4D data were deconvoluted using a constrained iterative and advanced maximum likelihood algorithm (iteration: 5), using Olympus Cellsens software (Olympus, Inc). Each 4D tiff stack file was processed using Fiji ^62^ (Figure S1). The deconvoluted images were processed using Gaussian blur (sigma = 2) and an image J plug-in “attenuation correction” (opening =3, reference =15) ^63^. A 10-µm W x 32 µm H boxed region corresponding to the contractile ring was selected and adjusted for the fluorescence intensity so that the signal would not be saturated in the next step, and rotated by 3D projection (Brightest point, interpolation on). After selecting a plane of en face ring view, the images were segmented and quantified using an Image J plug-in Morpholib J ^64^. The segmented contractile ring areas were measured for ring radius, ring centroid, and ring angle. The radius of the segmented area was estimated using ellipsoid fitting and derived by calculating the average of major and minor radii of the ellipsoid. Data were aligned relative to the time point first exceeded 10% ring closure. The ring trajectory images were obtained with an in-house image J macro using same data.

### Quantification of the intact AB contractile ring dynamics

Since the AB cell undergoes rotation during its division, the precise quantification is challenging. We first selected samples undergoing unilateral cleavage roughly in parallel to the imaging plane, and corrected for cell rotation using Stackreg plug-in of Image J ^65^, using chromosome signal (polar bodies were deleted using a brush tool to avoid abnormal image registration) (Figure S5). The rotation-corrected images were then processed by the same pipeline used for the P_0_ cell.

### 3D visualization and quantification of contractile ring dynamics in isolated blastomeres

The oblique view of cells in Figures 1A, 7A–B, and 7F were generated using an Image J plug-in Clear Volume ^66^. The analysis of the contractile ring dynamics in isolated blastomeres is challenging since the cell is rotated along different axes, and not all the planes were captured during imaging. Thus, we made en face ring view images for each time point using Clear Volume, and estimated the cell and ring contours, centroids, and diameters, by selecting more than four points along the cell and ring perimeter, respectively, using an Image J macro “Smallest Enclosing Circle” ^62^. The ring closure rate and eccentricity were calculated using the method described above for other cells. Measurements were performed three times per sample, and average values were used to mitigate the relatively higher error rate compared to the automatic segmentation method used for intact embryos.

### Kymographs

Kymographs in Figure 4D were generated by stacking 1 µm H x 40 µm W rectangular regions at a given time. Kymographs in Figure 4E were generated by stacking 11 µm W x 3.7 µm H rectangular regions corresponding to a myosin cluster. The timing is adjusted so that the peak myosin cluster intensity comes fourth out of a total of eight frames, using an in-house image J macro.

### Quantification of myosin enrichment rate

We measured myosin enrichment using the ring en face view of NMY-2::GFP, as shown in Figure 8B. Selection of the contractile ring region was performed using the segmentation data described above. However, unlike the highly pre-processed data used for segmentation (e.g., attenuation corrected), we used raw myosin data without deconvolution and other processes. The only processes performed were 3D projection and interpolation. After binarizing the segmentation data, we dilated the mask four times (mask 1) and also created another mask with four times erosion (mask 2). In our condition, mask1 covered most of the ring area, while mask 2 covered the area inside the ring (cytoplasm). The myosin signal at each time point was then quantified by subtracting the signal in mask 2 from the signal in mask 1. The timing of these time series data was aligned relative to 10% ring closure.

### Particle Image Velocimetry

A detailed image preprocessing pipeline is shown in Figure S3. Briefly, 4D data sets of NMY-2::GFP and GFP::-SAS-7-expressing embryos were processed with Gaussian Blur (sigma =1) and unsharp mask (radius =1, mask = 0.6) using Fiji. Centriole signals were deleted by filling zero values. After intensity adjustment to avoid signal saturation, half volume of image stack was filled with zero values to avoid projecting cortical myosin from opposite cortical sides. The images obtained were projected using an Image J function “3D projection.” We rotated 3D projected data relative to angle of cleavage at 25% closure and generated leading and lagging cortex views. Note that not always both lagging and leading cortices were clearly visible, so that in some embryos, either the leading or lagging cortex was used. Using these flattened cell cortex images, Particle Image Velocimetry was performed using MATLAB software PIV LAB (Algorithm: FFT window deformation, Interrogation area: 5.8 µm, Step size: 2.9 µm, Sub-pixel estimator: Gauss 2×3 point, Correlation robustness: Standard) ^43^. We rejected vectors exceeding 0.25 µm/s to remove estimation errors (the setting was visually confirmed to reflect the myosin foci movement). Obtained 3D matrix data (2D x time) were used for downstream analyses. The PIV matrix data were aligned relative to 10% ring closure, centered relative to the cell center corresponding to the furrow position, and resized to contain an entire embryo using Numpy and Pandas ^67, 68^.

### Estimation of cortical convergence

Cortical convergence was derived using the PIV vector matrix and a NumPy gradient function ^67^. When we define A-P-axis and circumferential-axis cortical flow velocities in an i x j matrix as u_i, j_ and v_i, j_, respectively, u_i+h, j_, u_i-h, j_, v_i, j+h_, v_i, j-h_ can be defined as follows using Taylor series ^69^:

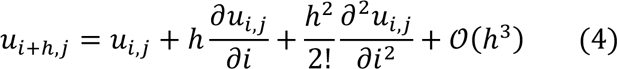

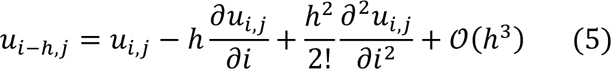

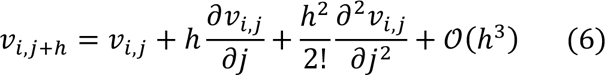

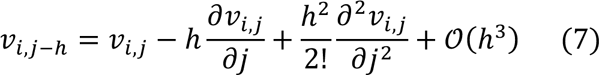

And we will obtain the following by subtraction:

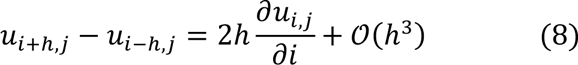

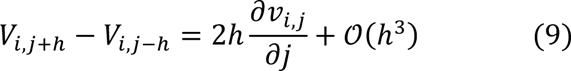

Thus, the cortical flow gradients at the interior points can be defined as follows:

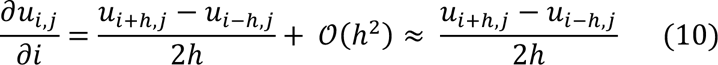

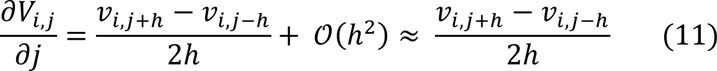

At the boundary, first order one-sided differences were achieved as follows:

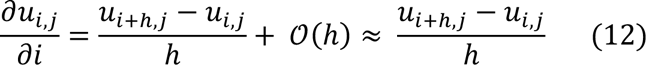

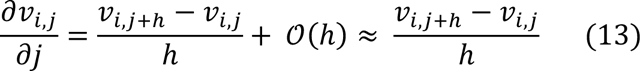

A-P-axis and circumferential-axis cortical convergence p_i,j_, and q_i, j_, respectively, were calculated as follows, where *a* is the PIV step size:

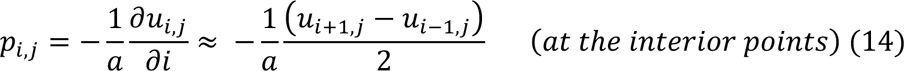

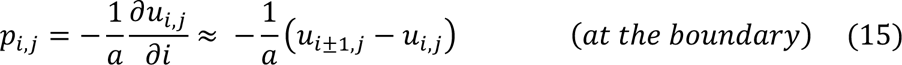

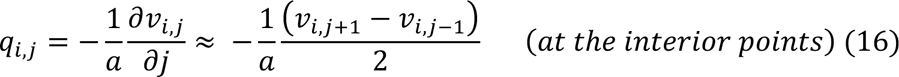

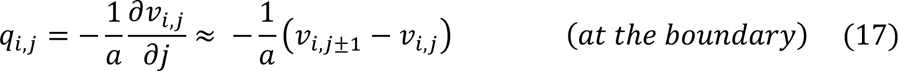

To quantify leading and lagging cortex convergence at the equatorial region, we calculated the mean convergence within a 6×3 grid (14.6 µm W x 5.9 µm H) around the center of the furrow position.

### Myosin foci tracking

Myosin foci were tracked using Manual Tracking plug-in of Image J.

In Figure 7D–E, cell position was registered using image J plug-in Stackreg, to eliminate the movement of myosin foci caused by the movement of cell body.

### Statistical analysis

For multiple comparisons, one-way ANOVA with the Holm–Sidak’s method was used. The comparison of two data was performed using the Welch’s t-test. No statistical method was used to predetermine the sample size. Error bars or bands correspond to the 95% confidence interval. The experiments were not randomized. The investigators were not blinded to the study. Statistical analyses were performed using either SciPy or Prism 9 (Graphpad). Symbols such as “****,” “***,” “**,” and “*” indicate *p* < 0.0001, *p* < 0.001, *p* < 0.01, and *p* < 0.05, respectively.

## Data availability

The data used in this study is available in the Source Data file.

## Code availability

The codes used in this study are available at https://github.com/sugi01/unilateralcytokinesis.

---

**Figure S1. Imaging and contractile ring analysis methods of P_0_ zygotes.**

(**A**) Imaging conditions and analytical flow charts. We used refractive index matching medium and polystyrene beads to prevent light scattering and cellular compression. The use of a refractive index matching medium reduced the required slice number from 34 to 31 and also improved image resolution, especially on the deeper side of the cell (focal planes far from the objective lens). (**B**) Temporal color-coded image of 20 frames around cytokinesis onset showing that compression-free embryos do not exhibit global cortical rotation. (**C**) Ring en face view images of 3D projected data and segmentation results. The first and last frames are the top left and bottom right in these figures, respectively. (**D**) Contractile ring closure curve after alignment of time series data relative to 10 % ring closure. Error bars indicate 95% confidence intervals. As we also measured myosin signal intensity and cortical flow dynamics using the same data set, all the data were aligned using the time relative to cytokinesis onset (or normalized time relative to cytokinesis onset) unless otherwise specified.

**Figure S2. Ring closure dynamics in RNAi and mutant backgrounds tested in this study.**

(**A**-**F**) Ring closure, Eccentricity, and Ring edge advancement curves were plotted against time relative to cytokinesis onset. See Figure S3 for the data plotted against normalized time. Lagging furrowing onset was defined as the time when the L_lag_(t) exceeded 0.02 (2% relative to initial ring size). (**G**) Relationship between peak eccentricity and time lag (raw values without normalization).

**Figure S3. Generation of cortical myosin data used for Particle Image Velocimetry (PIV) analysis.**

(**A**) An image preprocessing pipeline used to generate cortical myosin data for PIV analyses. See also Methods section. During the quality control step, very noisy or dim images were excluded from PIV analyses. These images are usually from image planes furthest from the objective lens.

**Figure S4. Cortical convergence in let-502(RNAi) and pod-1(RNAi).**

(**A**) A-P axis and circumferential axis convergece curves plotted against actual time (in seconds)

**Figure S5. Imaging and contractile ring analysis methods of the two-cell stage AB cell.**

(**A**) Imaging conditions and analytical flow charts. Intact AB cells were analyzed similar to the P0 cells with some modifications to correct for cellular rotation. For isolated AB blastomeres, we used normal growth medium and a different analytical pipeline to estimate ring size and position due to the movement and rotation of the cells. Isolated AB cells were measured three times, and average values were analyzed.

**Movie S1. Dynamics of cortical myosin and contractile ring of a single embryo**

Left top: maximum projection of cell surface NMY-2::GFP. Bottom left: ring en face view of NMY-2::GFP. Bottom middle: segmented ring. Bottom right: reconstructed ring trajectory. Right graph: Contractile ring dynamics of the same embryo. Note that the same embryo was used to generate all data.

**Movie S2. Cortical flow and flow vector map using Particle Image Velocimetry**

The cell surface NMY-2::GFP of leading (left), lateral (middle), and lagging cortex views (right). Yellow arrows are generated by Particle Image Velocimetry. Times are relative to cytokinesis onset.

**Movie S3. Dynamics of cortical compression and cortical ingression in *act-2(or295)***

Cell surface (top) and ring en face view (bottom left) of NMY-2::GFP. Ring outlines and centroid trajectory were color-coded as in Figure 6E (yellow: movement towards the the contraction site, blue: movement away from the relaxation site).

