## Supplementary material for "Contractile ring mechanosensation and its anillin-dependent tuning during early embryogenesis": Figure S1

### Figure S1 (Hsu, Sangha, Fan, et al.)

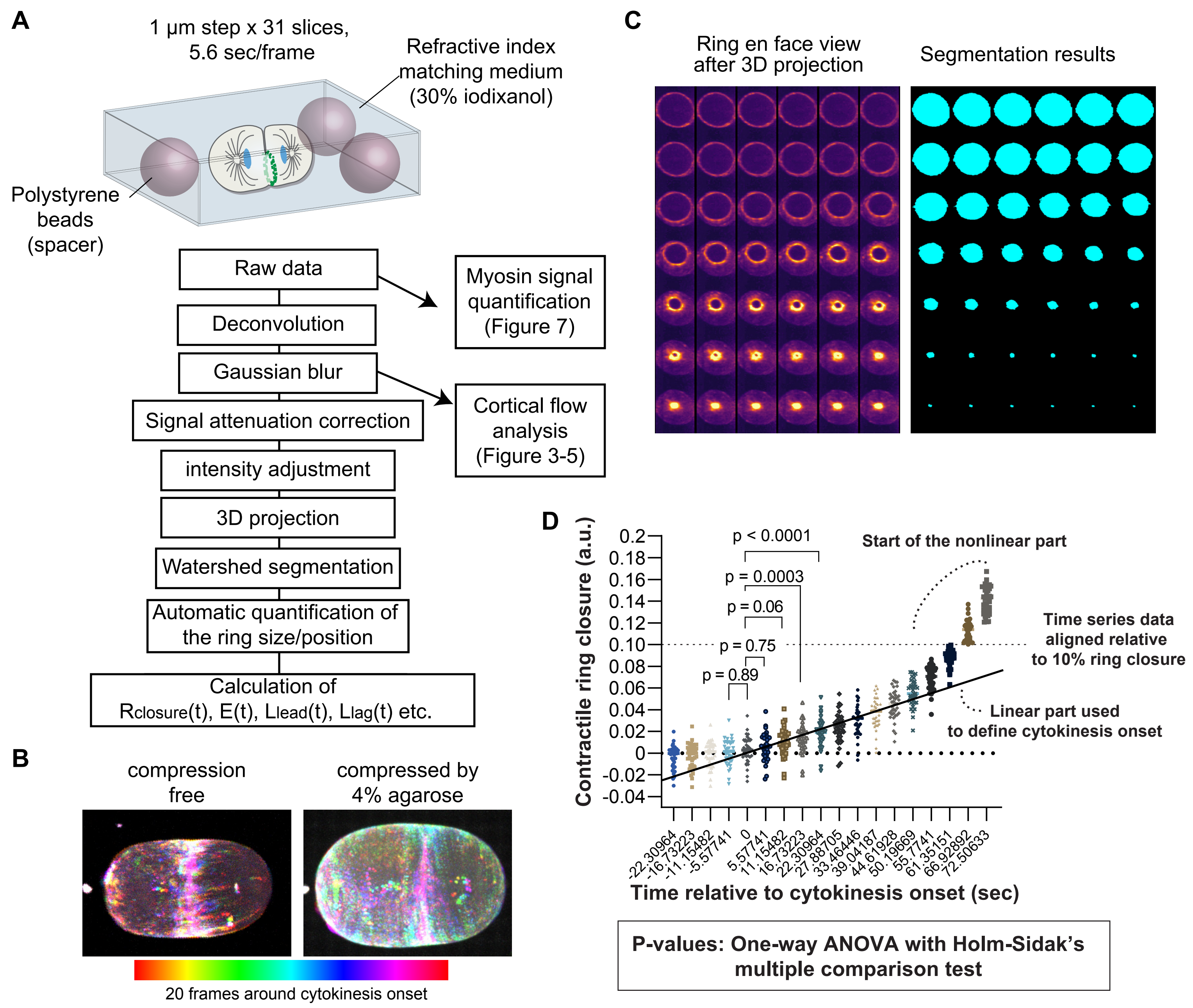


**Figure S1. Imaging and contractile ring analysis methods of $P_0$ zygotes.**
(**A**) Imaging conditions and analytical flow charts. We used refractive index matching medium and polystyrene beads to prevent light scattering and cellular compression. The use of refractive index matching medium reduced the required slice number from 34 to 31 and also improved image resolution especially in the deeper side of the cell (focal planes far from the objective lens). (**B**) Temporal color-coded image of 20 frames around cytokinesis onset showing that compression-free embryos do not exhibit global cortical rotation. (**C**) Ring en face view images of 3D projected data and segmentation results. The first and last frames are the top left and bottom right in these figures, respectively. (**D**) Contractile ring closure curve after alignment of time series data relative to 10 % ring closure. Error bars indicate 95% confidence intervals. Because we also measured myosin signal intensity and cortical flow dynamics using the same data set, all the data were aligned using the time relative to cytokinesis onset (or normalized time relative to cytokinesis onset) unless otherwise specified.
