## Supplementary material for "Contractile ring mechanosensation and its anillin-dependent tuning during early embryogenesis": Figure S2

### Figure S2 (Hsu, Sangha, Fan, et al.)

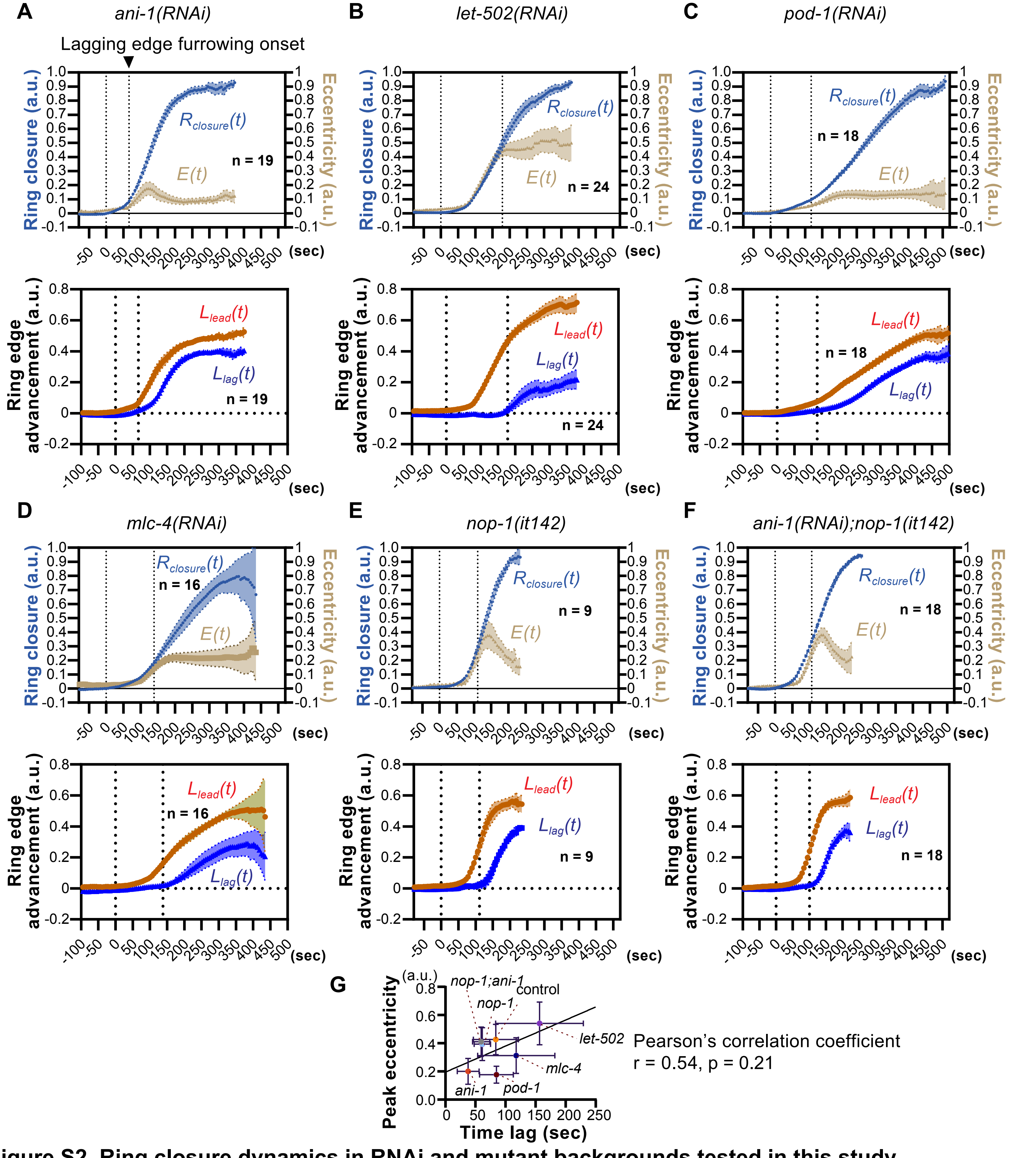


**Figure S2. Ring closure dynamics in RNAi and mutant backgrounds tested in this study.**
(**A-F**) Ring clousre, Eccentricity, and Ring edge advancement curves were plotted against time relative to cytokinesis onset. See Figure S3 for the data plotted against normalized time. Lagging furrowing onset was defined as the timing when the $L_{lag}(t)$ exceeded 0.02 (2% relative to initial ring size). (G) Relationship between peak eccentricity and time lag (raw values without normalization).
