## Supplementary material for "Contractile ring mechanosensation and its anillin-dependent tuning during early embryogenesis": Figure S3

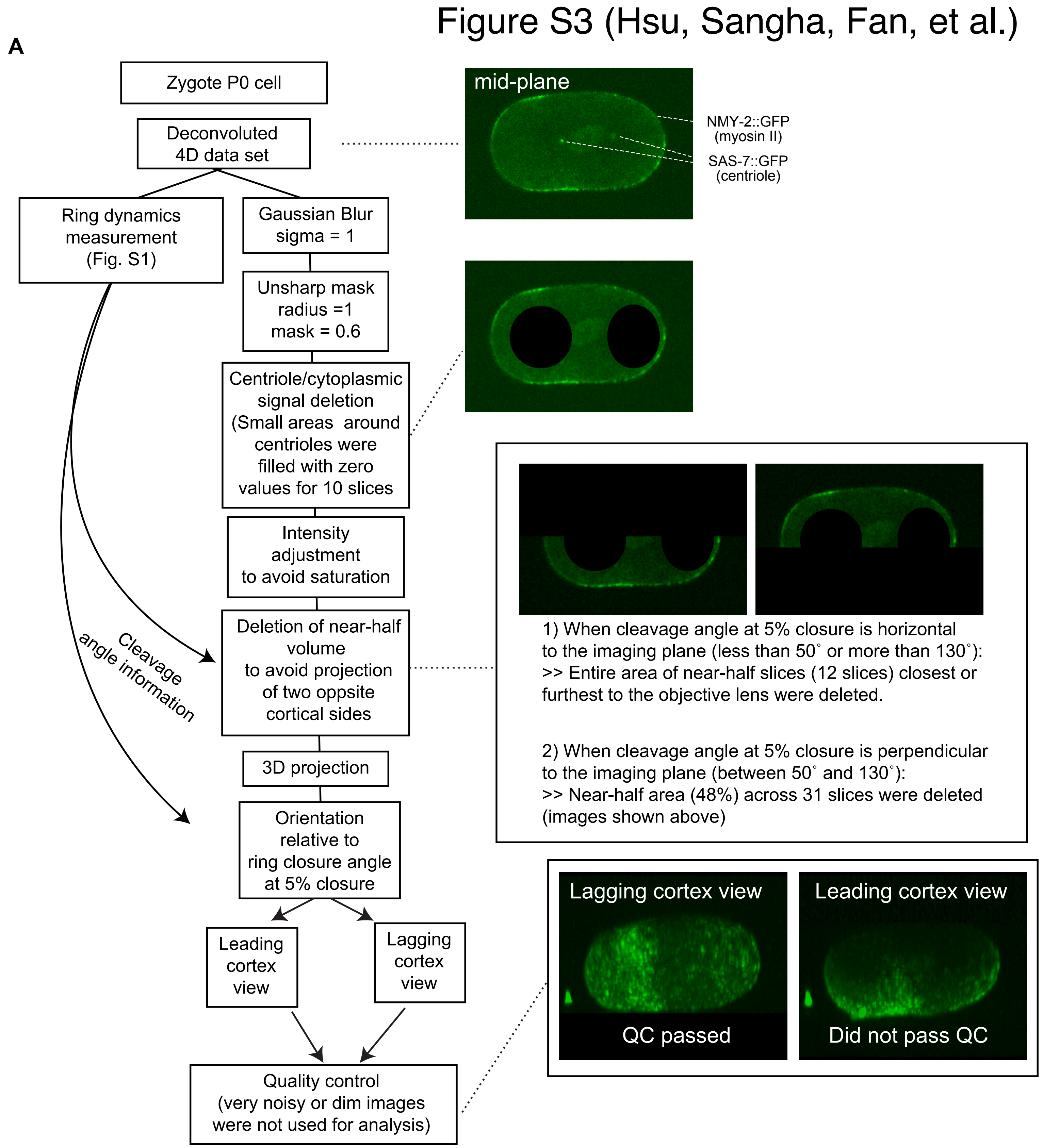


**Figure S3. Generation of cortical myosin data used for PIV analysis.**
(**A**) An image preprocessing pipeline used to generate cortical myosin data for Particle Image Velocimetry analyses. See also Methods section. At the quality control step, very noisy or dim images were not selected for PIV analyses. These images are usually from image planes furthest from the objective lens.
