## Supplementary material for "Contractile ring mechanosensation and its anillin-dependent tuning during early embryogenesis": Figure S4

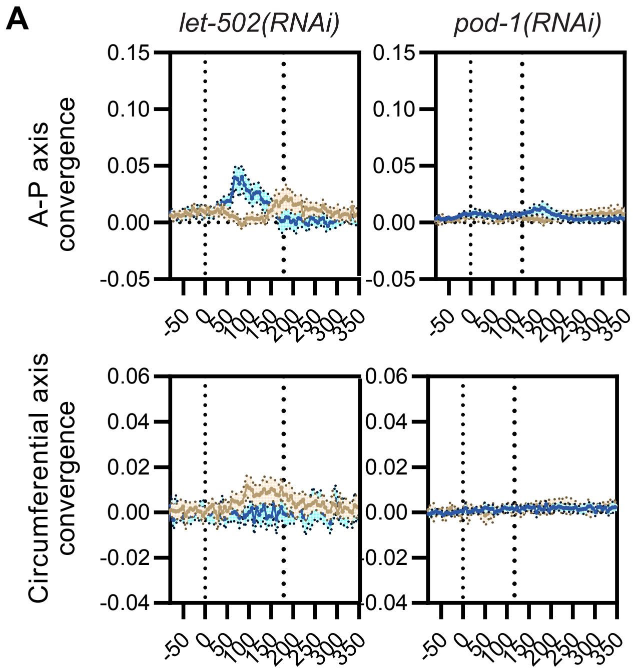


**Figure S4. Cortical convergence in *let-502(RNAi)* and *pod-1(RNAi).***
(**A**) A-P axis and circumferential axis convergece curves plotted against actual time (unit: sec)
