## Supplementary material for "Contractile ring mechanosensation and its anillin-dependent tuning during early embryogenesis": Figure S5

Figure S5 (Hsu, Sangha, Fan, et al.)

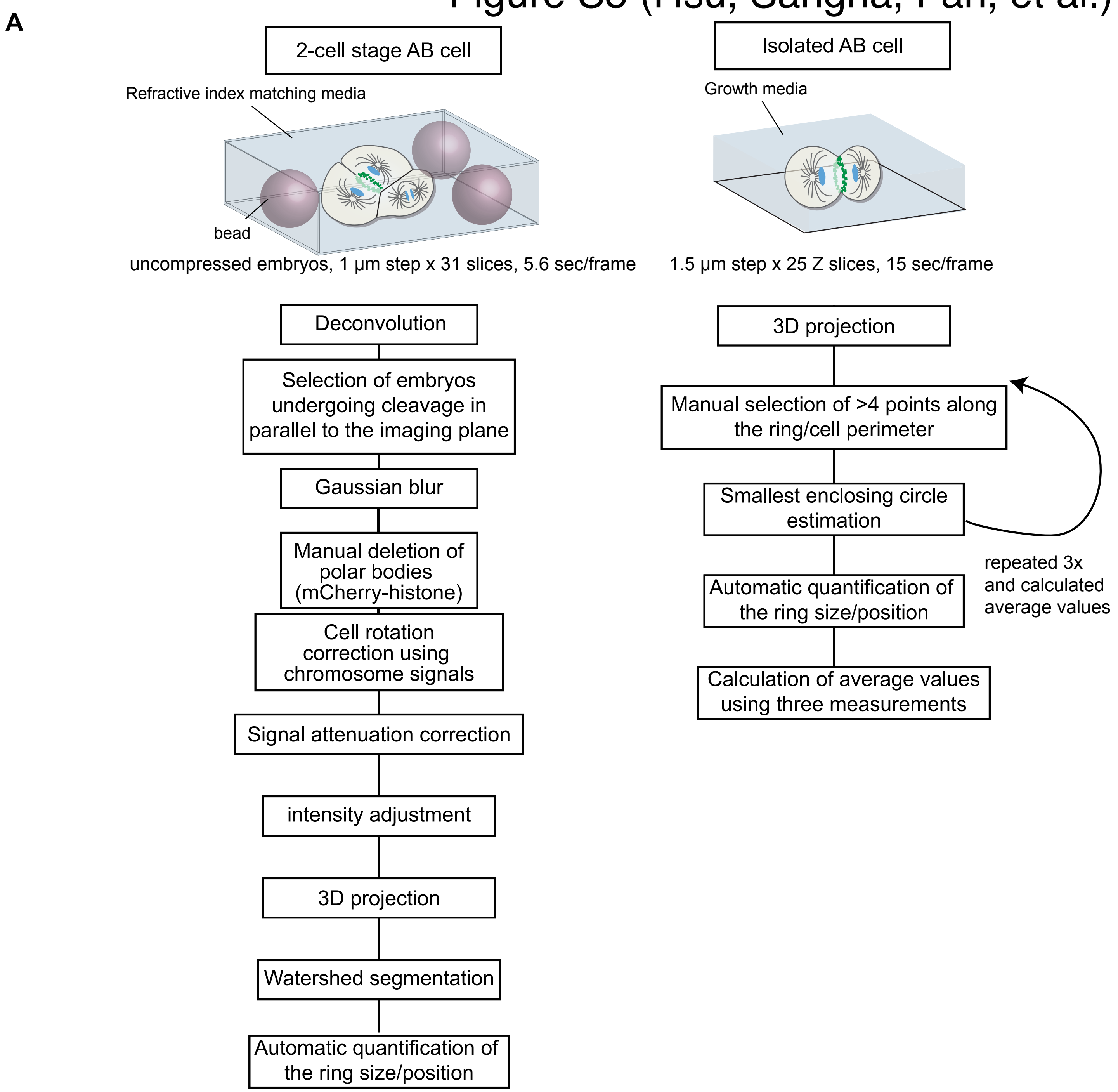


**Figure S5. Imaging and contractile ring analysis methods of the two-cell stage AB cell.**
(**A**) Imaging conditions and analytical flow charts. Intact AB cells were analyzed similar to the P0 cells with some modifications to correct for cellular rotation. For isolated AB blastomeres, we used a normal growth medium and a different analytical pipeline to estimate ring size and position because of the cells' movement and rotation. Isolated AB cells were measured three times, and average values were analyzed.
